# Death recognition by undertaker bees

**DOI:** 10.1101/2020.03.05.978262

**Authors:** Ping Wen

## Abstract

Dead conspecifics removal is important of being social to avoid pathogen transmission, which resulted in the evolution of a specific caste of undertaking workers in all hives bee species. However, it is mysterious that how the undertakers distinguish death and life instantly. Through integrative studies of behavioural tests and chemical analyses, a novel mechanism for dead conspecifics recognition is found in the Asian bee *Apis cerana cerana* Fabricius. The bees detect quickly the death of conspecifics based on decreased cuticular hydrocarbon (CHC) emissions, caused by the cooling of the dead bee. Specifically, with the decline of body temperature in death, the CHC emission was reduced. Undertakers perceived the major CHCs. Addition of synthetic CHCs, followed by heating, inhibited undertaking behaviour. Among these CHCs, heptacosane and nonacosane are the major compounds in a natural bee hive, providing a continuous signal associated with life. Via changing the vapour pressure then the ratio of emitted compounds encoding the physiological status of signal sender, insect chemical communication can be finely tuned by body temperature. This straightforward death recognition mechanism requiring little cost can be universal in animal living in social groups, especially in the social insects. Body temperature affected behaviour can response to increasing frequency of extreme weathers in global climate change, which help explain the recent worldwide bee health problem.

## Introduction

Recognition of death is an important behaviour of being social. In social groups, detection of death and rapid removal of corpses is essential in order to defend the colony against pathogen transmission (1–4). Humans use the cessation of heartbeats and breathing or cooling body temperature to diagnose death. All these even require complicated processing of information due to the development of resuscitation skills. However, what mechanisms are available to much more primitive organisms such as insects to assess whether conspecifics are alive or dead? It has been a long time, that carbonic acids derived from corpse decompose are responsible for the death recognition of dead bodies in social insects (2, 5). But this leads to theoretical and empirical conflicts. Theoretically, dead individuals, usually old or diseased workers, are at the highest risk of carrying pathogens. Leaving dead bodies not removed till decomposition increased the risk of pathogen transmission within the nest, which is conflicted with the importance of corpse management in social behaviour. In observations, specialized workers in eusocial species perform undertaking behaviour, removing the dead bodies of workers speedily (3, 6–8). Usually, a dead bee can be recognized within 30 min (7), before significant decomposition occurs with the concomitant production of odours of decay. In the darkness of a bee hive, no visual cues are available, and tactile cues from a freshly dead bee are probably indistinguishable from those of a living bee. Thus, how do undertaking workers recognize dead bees?

Chemical cues specific to life or death could signal rapidly death in social insects (3, 5). For example, in ants, it has been shown that the decrease in chemical signals is correlated with death. Two specific chemical compounds associated with life, dolichodial and iridomyrmecin, were identified in the Argentine ant, *Linepithema humile*, and these compounds inhibited necrophoresis behaviour (8). Also, the dead worker of termite *Reticulitermes flavipes* produce 3-octanone, 3-octanol as death signal (9). In honeybee, the blend of β-ocimene and oleic acid acts as a cue for diseased brood in hygienic behaviour of *Apis mellifera* (10). But it is not known if there is any compound responsible for recognizing the death of workers.

The dynamics of emission of a signal are related to the signal’s behavioural function in a social context. For a recruitment signal, release should be positively correlated with the number of individuals involved (11, 12). In contrast, a decrease in inhibitory signals can result in the release of an inhibited behaviour. Based on the correlation between the dynamics of signal emission and the behavioural process being affected, signals mediating certain behaviours can be identified, for example in the identification of recruitment signals in termite foraging (13), honeybee alarm communication (14) and most sex pheromone in calling moth (15). In contrast to the collective decision making by a large number of workers in swarming behaviour (16, 17), undertaking bees are a small subset of individuals in the hive. The behavioural process of undertaking does not require the involvement of large numbers of individuals, so recruitment signal is not likely to exist for the recognition of dead bees. Also, undertaking bees can be experienced in the removal process (18), but there is no appetitive reward or aversive punishment to undertaking bees that may generate the associative learning of olfactory cues. Thus, the perception of chemical cues associated with life and death is likely to be innate (7).

Cuticular hydrocarbons (CHCs) have multiple functions in social insect communication (19, 20). They typically consist of long chain hydrocarbons with more than 20 carbons, characterized by low vapour pressure, and function as low-volatility signals in short distance or contact communication (21). Most known queen pheromones or reproductive-specific signals in wasps, bumble bees, ants, and termites are CHCs (22–24). Even behavioural polymorphism within a caste is regulated by CHCs, for example in *Pogonomyrmex barbatus* (20). Besides the Nasonov pheromone, CHCs are the major cuticular compounds in honeybee (25), functioned as colony discrimination signals (26–30) and even behavioural polymorphism signals (31). In waggle dance communication, the detection of CHCs can stimulate followers (32). Besides signalling, CHCs have additional roles in providing physical protection of the colony. For example, CHCs are major components in the wax comb as building materials (33). Also, CHCs are abundant on the body surface of most insects, serving to maintain water balance. The persistence of waxy CHCs due to their low vapour pressure and stability is suited to these physiological functions (34). Individuals thus need to balance the two roles of releasing CHCs for short distance communication, while also producing and retaining CHCs in larger quantities for desiccation resistance. In honeybee, besides contact cues, short distance communication can be signalled by semivolatile CHCs; for example, CHC headspace volatiles detected in the waggle dance area are different from those in other areas of the nest (32).

Social insects can discriminate CHCs. In *Camponotus* ants, single sensillum recording (SSR) and learning and memory tests showed that a broad spectrum of CHCs can be detected by the olfactory neurons but with precise discrimination (35). Honeybees can also perceive and discriminate among different CHCs in learning and memory assays (36, 37). But it was not directly shown if honeybees can discriminate CHCs physiologically or not.

Living honeybees have body temperatures ranging from 35 °C to 45 °C (38–40), usually higher than the hive wall temperature under ambient conditions. When a bee dies, the decrease in body temperature reduces the vapour pressure of its CHCs, decreasing their emission. This might have helped the chemical signalling in nestmate recognition (38). Thus, rather than using decreasing body temperature as a death signal, honeybees may perceive the reduction of CHCs in the headspace around a dead individual as a death signal. Along these lines, Visscher (1983) showed that addition of paraffin (containing many of the alkanes found in honeybee CHCs) inhibited necrophoresis behaviour in *Apis mellifera*, and exhaustive extraction by Soxhlet extraction also lead to failure of death recognition (7). With increasing frequency of extreme weather events in recent years (41), any temperature based behaviour can response to climate changes. But till now it is not known how bee health was affected by climate change directly.

Till now, no chemical life and/or death signal was identified in honeybees, and little on this was known in most social insects. By knowing chemical cues for death recognition, we can assess the complexity and diversity of this widely distributed behaviour in social insects, which helps explain why social behaviour can be evolved multiple times from different lineages. Here, my objective was to identify the signals used to detect death in the Asian honeybee, *Apis cerana*. By using headspace volatile collection, gas chromatography (GC), and coupled gas chromatography-mass spectrometry (GC-MS), I compared the volatiles from living and dead bees. I analysed the effect of body temperature of living bees on the evaporation of CHCs using thermal imaging and simulation. I tested the antennal perception of bees to specific CHCs, using coupled GC-electroantennography (GC-EAD). Then I performed reduction and addition bioassays with CHCs, testing the hypothesis that the decrease of CHC emissions signalled the reduction of body temperature as a specific signal eliciting honeybee undertaking behaviour.

## Results

In performing undertaking behaviours, undertaking bees would palpate a dead bee, check around it several times and then directly grab the dead bee and attempt to fly away. Dead bees were dropped at distances away from the colony in observations. Basically, the dead bees were recognised by their reduced cuticular hydrocarbon emissions (Fig 1.).

**Figure 1.**
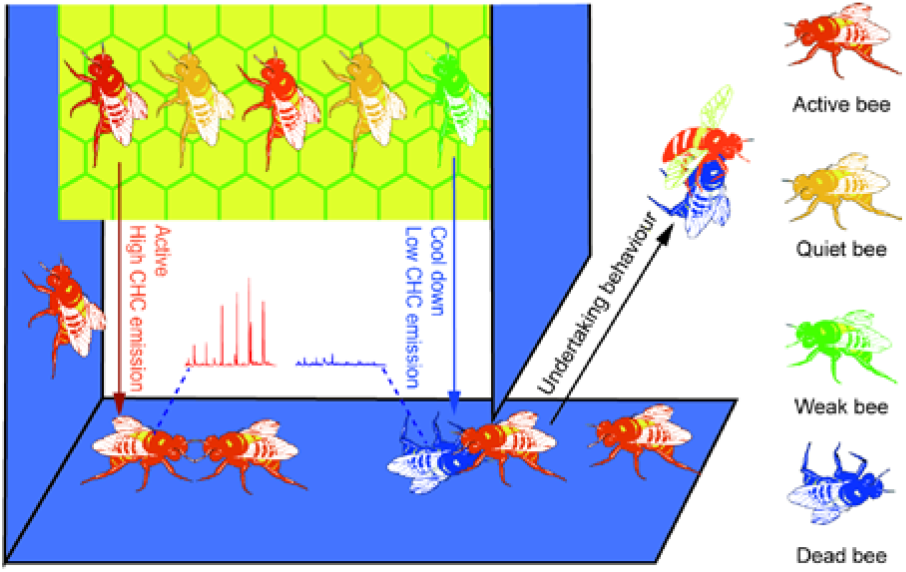
Undertaking behaviour is triggered by low CHC emission. Bees are coloured according to their status and body temperature as shown in the right side. Active bees were warm in red, while dead bees were cool in blue. Hive parts are also coloured in the similar pattern. The comb is warm in yellow, while the hive walls are cool in blue.

### Life signal is cuticular

Bioassays of worker body sections showed that body parts were removed selectively (GLMM: *G^2^_7_* = 56.375, *P* < 0.001); the more cuticle on the body parts, the more they were removed (*Sig* = *P* < 0.05). The stings with alarm pheromone as controls were always ignored (*P* = 0.74 > 0.05). Despite the abundant volatiles emitted as alarm pheromones, the stings are not removed more than clean filter paper controls, which excluded the possibility of using alarm pheromones as a signal of life (Fig. 2A).

**Figure 2.**
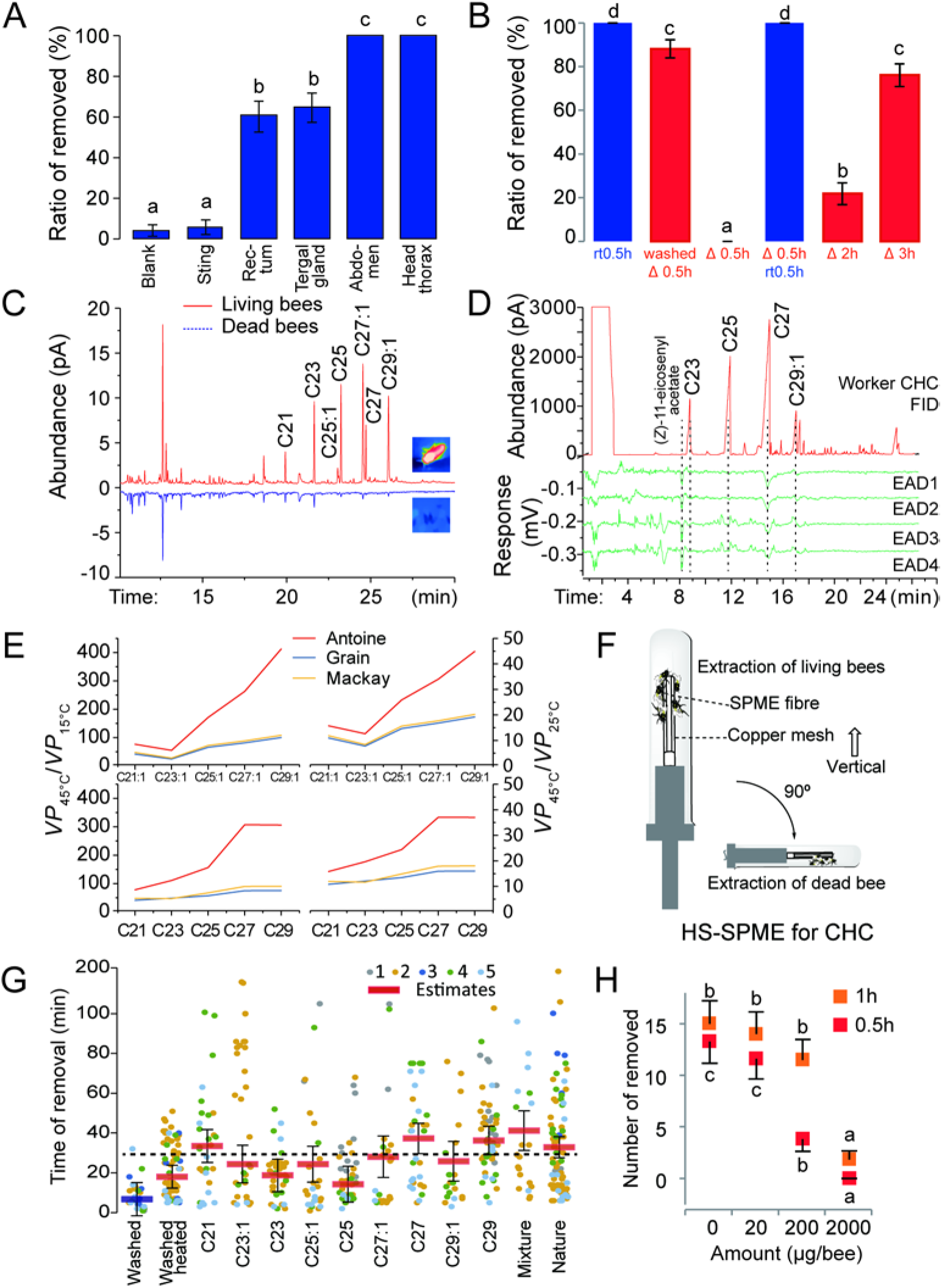
Honeybees perceive death from the reduction of CHC emission. A) Localization of death recognition signals to the cuticle. Blank: a filter paper without any body parts; Abdomen: the abdominal cuticle. Columns with different letters were significantly different at the *P* < 0.01 level. B) Effect of heating on undertaking behaviour. Column and samples are coloured in red for heated (Δ) and in blue for ambient temperature (rt). Columns with different letters are significantly different at the *P* < 0.01 level. C) Comparative analyses of the HS-SPME extraction of cuticular volatiles between living and dead bees. D) GC-EAD analysis for the perception of CHCs. E) Changes of CHC vapour pressure at different body temperatures of living (45 °C) and dead bees (15 °C ~ 25 °C). F) Diagram for HS-SPME collection of bee volatiles at close range. G) Undertaking inhibition bioassay for CHC addition and reduction on dead bees. Columns higher than the dashed line are significantly different (*Sig = P* < 0.05) from the washed bees tested at ambient temperature as control. Mixture is the mixture of synthetic CHCs as described in methods. Nature is heated dead bee. Different colours of points represent different colonies. Columns are coloured in red for heated and in blue for ambient temperature. H) Undertaking inhibition bioassay by adding wax solution to dead bees. Columns marked with different letters are significantly different (*Sig = P* < 0.05). Data recorded successively at 0.5 h (error bar downward) and 1 h (error bar upward) were analysed separately but plotted together.

### Effect of heating and/or washing on undertaking

Active living bees on the floor area had an average body temperature of 43.6 ± 1.7 °C (mean ± SD, *t-test*: *t* = 0.018, *df* = 164, *P* = 0.99), whereas dead bees cooled to ambient temperature (15 °C to 25 °C during tests) soon after perishing. By using 3 equal sized colonies, no effect from colony (GLMM: *G^2^_2_* = 0.00, *P* = 1.0) or colony*dead bee treatments interaction (GLMM: *G^2^_10_*=4.11, *P*=0.942) was observed on the ratio of dead bees removed. The effect of dead bee treatments with heating, cooling and body wash at different times on the ratio of dead bees removed was significant (GLMM: *G^2^_5_* = 65.04, *P* < 0.001). Fresh dead bees were all rapidly removed under natural ambient temperatures in 0.5 h. In contrast, heated dead bees were removed only after much longer periods (up to 2 h) or if they were allowed to cool to ambient temperature (Fig. 2B). Also, washed bees deprived of most of their CHCs and heated were always removed (EM: 88.0 ± 4.2 %, n = 72) more than heated unwashed dead bees (EM: 0.0 ± 0.0 %, n = 72, *P* < 0.001), but as much as long-time (3h) heated dead bees (EM: 76.0 ± 5.2 %, n = 72 *P* = 0.144 > 0.05) (Fig. 2B). These two results suggested that cuticular signals affected by body temperature might be regulating undertaking behaviour.

### CHC emission diminished on dead bees

HS-SPME-GC analysis revealed that CHC emissions decreased after death in *Apis cerana.* Twelve peaks were significantly higher in the headspace volatiles of living bees than in dead bees (paired t-test: *n* = 9, *sig* = *P* < 0.05) (Fig. 2C). Nine of these compounds were CHCs, including heneicosane (C21) (*t* = 9.99, *df* = 8, *P* < 0.001), tricosane (C23) (*t* = 11.28, *df* = 8, *P* < 0.001), (Z)-7-pentacosene (C25:1) (*t* = 2.74, *df* = 8, *P* = 0.026), (Z)-9-pentacosene (C25:1) (*t* = 4.80, *df* = 8, *P* = 0.001), pentacosane (C25) (*t* = 4.88, *df* = 8, *P* = 0.001), (Z)-9-heptacosene (C27:1) (*t* = 3.20, *df* = 8, *P* = 0.013), heptacosane (C27) (*t* = 4.76, *df* = 8, *P* = 0.001), nonacosenes (C29:1) (*t* = 4.04, *df* = 8, *P* = 0.004) and nonacosane (C29) (*t* = 3.37, *df* = 8, *P* = 0.010). GC-EAD analyses showed that undertaking bees could detect all the major CHCs, including C23, C25, C27, and C29, and (*Z*)-11-eicosenyl acetate, a contaminant from venom (Fig. 2D). Among the four CHCs, C27 and C29 generated the largest responses.

### Vapour pressure linked body temperature and CHC emission

Vapour pressure calculations showed that within the body temperature changes, C27:1, C27, C29:1 and C29 had the biggest changes in vapour pressure (Fig. 2E). The vapour pressure controlled the evaporation, and consequently the quantities of CHCs in the headspace volatiles. Thus, the evaporated CHCs, especially C27 and C29, were reduced substantially between live and dead bees.

### Synthetic CHCs inhibits undertaking under active body temperature

In assays of the synthetic CHCs, undertaking bees checked the CHC-modified dead bees continuously until they decided to carry the dead bees away. From the durations recorded, there were significant effects from the colony (GLM: *F_(4,362)_* = 15.50, *P* < 0.001), the environmental temperature (*F_(13,362)_* = 2.03, *P* = 0.018 < 0.05), the tested samples (*F_(12,362)_* = 6.50, *P* < 0.001), the interactions of colony*samples (*F_(19,362)_* = 2.58, *P* < 0.001) and the interaction of temperature*samples (*F_(50,362)_* = 1.94, *P* < 0.001). Generally, the larger the colony, the faster they removed the dead bees (*P* < 0.05), and the undertaking behaviour was inhibited best when ambient temperature was 25 °C (*P* < 0.05). Extracted dead bees tested at ambient temperature were removed quickly (EM: 6.7 ± 4.25 min, n = 26). Addition of C21 (EM: 33.5 ± 4.21 min, n = 30), C27 (EM: 37.3 ± 3.88 min, n = 33), C29 (EM: 36.3 ± 3.64 min, n = 52), and a mixture of CHCs (EM: 41.2 ± 5.06 min, n = 20) to extracted and heated bees significantly inhibited undertaking behaviour (*P* < 0.05) (Fig. 2G). Also, treatment of dead bees with wax solution also inhibited undertaking, with a significant effect of the quantities of wax added to the number of dead bees removed (GLMM: *G^2^_2_* = 12.723, *P* = 0.002 < 0.05). The more wax added, the stronger the inhibition (*P* < 0.05).

## Discussion

### Body temperature-vapour pressure correlation in bee CHC emissions

Results suggest that the bee death is encoded by the reduction of CHC emissions caused by the cooling of a dead bee on the hive floor. Undertakers apparently do not use the decrease of body temperature as a life signal (see Fig 2 B,G Washed but heated bodies were removed quickly.), even though they have thermo receptors (42) and may use it in nestmate recognition (38). Instead, undertakers use the associated reduction of CHC volatiles as an indication of death. On dead bees, the CHCs retained on the body surface as contact cues do not change within the short time when undertaking behaviour occurs (Fig. S3). Undertaking bees appear to use the decrease of headspace CHC volatiles (emitted CHCs) to discriminate living or dead bees. Due to the low vapour pressure of CHCs, researches on CHCs in contact communication has used solvent extraction or SPME fibre wiping to sample CHCs on the body surface, but not the actual amount of CHCs in the headspace that may function in short range communication. Short-distance headspace volatile collection can explain more precisely the role of CHCs emission in insect communication.

Ratios of signal compounds encode behavioural specific information in social insects (11, 13). Although SPME is not accurate in detecting the actual ratios of CHC compounds, basically, the ratio of compounds in the volatiles is quite different from the ratio of compounds in the whole body extract under the same HS-SPME extraction method (Fig. 2C, D). In headspace samples, C27:1 and C29:1 were the major compounds, whereas in whole body extracts, the saturated alkanes C23, C25, C27, and C29 were the major compounds. The calculation of vapour pressures predicted that the volatilities of C27:1, C27, C29:1 and C29 would be most affected by variations in body temperature. This could explain the ratio differences, but I cannot exclude the possibility that C27:1 and C29:1 in the headspace volatiles were from glands or internal tissues with smaller cuticular area (43). The relatively higher vapour pressure of C21, makes it the major emitted CHC in the headspace around a bee, although its quantities on body surfaces were relatively low compared to other compounds. Coincidentally, C21 is the reproductive-specific signal in the termite *Reticulitermes flavipes* (23). In calculations, the alkenes C27:1 and C29:1 showed the biggest changes in volatility with body temperature, but they were not significantly active as signals of life. Instead, bees attracted to the artificially warmed dead bees with added C27:1 and C29:1 samples performed balling behaviour (44). However, these compounds are only present in trace amounts on the body surface (See FID of whole body extract in Fig. 2D), so it is unlikely that they would be reliable signals of life. Further tests are needed to show whether these compounds are involved in defensive behaviours of honeybees.

Body temperature of active insects are always higher than environmental temperature, for example in singing bladder cicadas (45), flying sphinx moth (46). Body temperature variations between physiological status such as active and quiescent conditions are sure to affect greatly the vapour pressure of pheromone compounds secreted from body surface. And it is common for insect to use hydrocarbons of different molecule size for communication (15). Major CHCs studied here were differed by 2 carbons between, resulting in highly varied ratios of the compounds emitted according to the changing body temperature under different physiological status (see calculation results in Fig. S4). Since ratios of compounds are always encoding species-specific or physiological and behavioural specific information, by considering the body temperature affected ratio of volatiles, more details can be explicated in insect chemical communication, especially in the social insect communication, which is mediated mostly by CHCs.

### Precise perception to CHCs in communication

In GC-EAD tests, all CHCs lacking an oxygen-containing group elicited weak responses (47, 48). Thus, EAD systems with a high sensitivity amplifier (14, 49) and static noise control (filtering the air flow with copper mesh and a stainless steel shield cage) were used in this research. The setup enabled the first EAD recording of honeybee responses to CHCs. The biased EAD responses to different CHCs indicated the discrimination ability of olfactory receptors. This can be further investigated with SSR tests. Moreover, processing of signals in the brain, leading to behavioural outputs, can be controlled by neural hormones (50), so that the peripheral perception of compounds may not determine the behavioural responses. Thus, comparative recordings of neural responses to CHCs in undertakers, foragers, and guards may help explain the behavioural polymorphism in honeybee workers. Based on clear discrimination of compounds, with C23:1, C23, and C25 encouraging waggle dance followers in *Apis mellifera* (32), and C21, C27, and C29 encoding life status in *A. cerana* found here, the differentiation of the behavioural function of CHC indicates again the complex chemical communication system in honeybee hives that is still relatively poorly understood.

By considering the effect of body temperature, CHC-based honeybee communication can be explained in more detail. In the complex background odour, the use of variable ratios of headspace CHCs could alleviate signal interference for more precise communication. Using thermal images of hives, detailed maps of CHC signalling can help explain much clearer the communication in various behaviours.

### Bee health response to climate change via affected body temperature

Here, temperature dropping scale affects the change of CHC emissions, allowing for recognition of dead bees and triggering of undertaking behaviours. Extreme weather conditions can affect the dropping scale of body temperature from alive to dead and weak bees in the hive. With increasing frequency of extreme weather events in recent years (41), the pathogen defence by undertaking and hygienic behaviour can be affected. Using the decrease of CHC signals to trigger undertaking behaviour can be inclusive. In extremely cold days, the cold anesthetized and/or weak bees may also be recognized and removed by undertakers (51) in the mornings. If not warm up, those cold anesthetized bees taken out of the hive will die. Also, in extremely hot days, hives may be subjected to high temperatures more often than previously, possibly increasing the chance of pathogen transmission through dead bee contact, via failure to recognize the dead or weak bees warmed up unusually. Here, together with the fact that temperature experience of brood affect the behaviour of adult bees (52), honey bees may response to the increasing frequency of extreme weather with high sensitivity, resulted in colony collapse disorder.

For beekeeping practice, with honeybee pathogens such as *Nosema* (53) and *Varroa* mite transmitted viruses (54) spreading worldwide, causing great losses in global apiculture, the optimization of undertaking and hygienic behaviour can help improve the resistance to pathogens. Understanding the chemical communication in undertaking behaviour provides a basis for developing methods for optimizing honeybee undertaking and hygienic behaviour. Thus, lowering the hive floor temperatures, for example, by adding a ventilated bottom panel to hives, may help the recognition of infected bees based on this research.

### Straightforward death diagnosis may help the evolution of sociality

Along the line of related researches (5), no compound of unique pheromone structure is needed for instant death recognition. Also, the CHCs signal for life in honeybees can be of same as CHCs used by solitary species in stress resistance, parasitism, aggression, competition and/or predation (34). Compared with the ants and termites using specific more volatile life signal (5, 8), the body size may affect the strategies used. The climbing tiny ants and termites are less likely to be warmed up to a much higher temperature than the ambient condition, which limited the chance of using CHCs for multi-purpose. But the ant and termite life and death compounds were also used in other behaviours. Thus, specifity of life or death signal can be simply quantitative not necessarily qualitative. Undertaking or necrophoresis, the important step of pathogen defence, is crucial for the evolution of higher sociality. Hereby, social species should spend neither more energy in evolving immunity with undertaking against pathogens transmission, nor more energy in evolving a new life signal for undertaking than solitary species. Simply, straightforward death recognition makes no more difficulty to become social safely.

## Supplementary information

Material and methods and supplementary figures and table were provided in Supplementary information. Crude data for figures were provided in the ZIP files.

## Acknowledgements

I thank Prof. Jocelyn G. Millar in UCR, Prof. Jin Chen and Prof. Zhanqi Chen in XTBG and Prof. James Nieh in UCSD for critic comments and edits. I am grateful to Lu Zi-Yun, Yang Zhen-Xin, Na Bo, Klett Katrina and Zhao Qin, for their help in bioassays. I thank Lin Hua in XTBG for kindly providing infrared camera. I acknowledge the Public Technology Service Centre of XTBG for providing help on instruments.

## Supplementary Information

### Materials and methods

#### Bees

The five *A. cerana cerana* Fabricius colonies used in this experiment were maintained at the Southwest Biodiversity Research Centre (SBRC, 25°08’21.2”N, 102°44’22.4”E) in Kunming, Yunnan, using standard beekeeping protocols. Natural nectar and pollen resources were available in the local habitat through the year. Observation hives were set up in a working shed at the same place. Workers which picked up dead bees were classified as undertaking bees. The five colonies were not always uniform in size, having varied number of individuals (Table S1).

#### Localization of death recognition signal

Foragers collected in front of the hive were killed by freezing at −40 °C for 10 min. These fresh dead bees were then dissected into different parts, including the rectum, the sting apparatus, the tergal glands, the head and thorax and the remaining abdominal cuticle. All these body parts were subjected to bioassay within 10 min after dissection. In bioassays, each body part was positioned on a clean filter paper disk (1.0 cm diameter, cleaned by soaking in pure water). To emit odour from internal tissue or glands, the head with mandibular glands and the rectum were squashed on the filter papers. All samples were positioned on the floor of observation hives. Tests were done under ambient hive conditions. Bioassays were carried out on sunny autumn days when colonies were relatively inactive. In 2 h, the number of samples removed and the total number of samples tested were recorded as event/trail data. Samples selectively picked up and then dumped outside the hive were considered to contain the signal for dead body discrimination.

#### Living bee body temperature measuring

To confirm the difference between the temperature of living and dead bees, a T1040 infrared camera (FLIR, Oregon, USA) was used to record high definition thermal images and video at the bottom of an observation hive. In this floor area where dead bees drop, there were usually moving living bees as well, often foragers coming in and out. The temperature of living bees on the recorded hive floor area was measured using FLIR tools software (FLIR, Oregon, USA).

#### Inhibitory heating bioassays

Because the reduction in headspace CHCs was recognized by undertaking bees, the effect of reducing body temperature on CHC emission was tested. I placed fresh dead bees as prepared above for whole body bioassays on two different glass Petri dishes (9 cm id) (Fig. S1), with one held at ambient temperature, and the other heated using a custom-made controller, composed of a 9 cm glass Petri dish, a W1209 thermostatic controller (Telesky, Shenzhen, CN), a 10 K MF58 thermistor (Risym, Shenzhen, CN) and a 15 W Cr20Ni80 heating wire (Yusheng, Jiangsu, CN). After calibration using the T1040 infrared camera (Fig. S1), in the dish, bees were heated to a steady temperature, approximating that of a living bee (38~41 °C). Temperatures inside and outside the hive were measured using a type K thermocouple read by a VC97 digital multimeter (Victor, Shanghai, CN). In the tested hives, the difference in average body temperature between a dead bee and a living bee was 10~25 °C. In treatments, dead bees which were not removed were considered as inhibiting undertaking behaviour.

#### CHC reduction and addition bioassays

Because the assays above suggested that CHCs were associated with the death signal, the effects of reduction and addition of CHCs were tested. Bees were freeze-killed as stated above, and the stings and venom sacs were removed so as not to agitate the bees in the hive. I removed the stings because the experiments above had shown that they were not active in necrophoresis, but would activate the hive, hampering undertaking behaviour. In CHC reductions, fresh dead bees were washed three times with 3 × 0.5 mL hexane in a 1.5 mL glass pipette, and then were taken out and dried for 10 min to evaporate the solvent. By washing with a small volume of solvent instead of Soxhlet extraction, the CHCs on the surface of the dead bee were not eliminated, but only reduced to about one fifth of original amount according to the GC analyses of the concentrated first and second washes of the same bees (Fig. S2). In CHC addition, three types of sample were prepared: (I) all single synthetic compounds were dissolved in hexane at 10 mg/mL; (II) for three isomers of each alkene, each isomer was added respectively at 6.6 μL (i.e., 66 μg/bee); (III) to simulate the natural CHC profile, a mixture of 8 mg C21, 3 mg C23:1s (Z5:Z7:Z9=1:1:1), 60 mg C23, 3 mg C25:1s (Z5:Z7:Z9=1:1:1), 70 mg C25, 3 mg C27:1s (Z5:Z7:Z9=1:1:1), 90 mg C27, 3 mg C29:1s (Z5:Z7:Z9=1:1:1), and 60 mg C29 were dissolved in 10 mL hexane. Extracted bees were dosed with 20 μL of these CHC solutions (i.e., 200 μg/bee). After addition, bees were dried and warmed for 5 min at 40 °C to evaporate the solvent, then introduced to the floor of the beehive. Bees were tested in groups of 3 to 6 per trial based on the activity of the hive (estimated by number of bees collected within 5 min at the hive entrance). Three extracted dead bees under ambient conditions and three extracted and heated dead bees were used as controls in all tests. The time between the test sample introduction and removal was recorded for each test bee. The numbers of replicates were decided by the numbers of workers that can be collected within 5 min in front of each tested hive on each day, which is affected by the size of the hive.

Major compound of bee wax are esters at 71%, the major one is triacontanyl palmitate (C46H92O2). Others are alcohol and acids. The ratio varies according to origin and species. CHCs are ~14% that can be volatile in normal temperature, other major compounds are esters with 30 more carbons that is not volatile (1,2). In another experiment, comb wax containing all the major CHC components was dissolved in hexane at 1 mg/ml, 10 mg/mL, or 100 mg/mL. Fresh dead bees were first washed as described above. Then the bees were dosed with 20 μL of the wax solutions, to make wax-coated dead bees with 20 μg, 200 μg, or 2000 μg per bee. The wax-coated bees were dried and heated as described above to remove solvent. Both fresh CHC reduced and added bees were placed in a 9 cm petri dish in the bee hive for undertaking bioassays. Each wax dose was replicated 15 times. Three colonies were tested. Natural dead bees were used as control. The numbers of removed and tested bees were recorded for each concentration in each colony.

#### Collection of volatiles from living and dead bees

To analyse the difference in CHC emissions between living and dead bees, the CHCs evaporated from bodies over short distances were compared using headspace solid phase micro-extraction (HS-SPME). Living bees were collected from three colonies. In each colony, 5 bees were collected without disturbance and then held in the dark in a clean glass test tube (18 mm id, 180 mm long). A 65 μm PDMS/DVB SPME fibre (Supelco, CA) protected by a copper mesh tube (40 mesh, 3 mm id, 50 mm long) was used to collect volatiles emitted by the bees. When extracting living bees, the apparatus was kept upright. Bees would keep moving upward to gather near the SPME fibre. The living bees were extracted in darkness first. I collected headspace volatiles from the living bee in the dark to reduce bee movement. Then, these bees were frozen and killed; their stings were removed using a pair of forceps as described above for the fresh dead bee samples. The dead bees were then placed near the SPME fibre. All the samples were extracted for 0.5 h under ambient conditions, with bees contacting the copper mesh but not the SPME fibre, to obtain the cuticular volatiles.

#### Chemical analysis

##### Chemical standards

Commercially available heneicosane, (*Z*)-9-tricosene (97%, in GC-FID peak area hence force), tricosane (99%), and pentacosane (99%) were purchased from TCI (City, Japan). Other CHCs, including (*Z*)-5-tricosene (98%), (*Z*)-7-tricosene (97%), (*Z*)-5-pentacosene (97%), (*Z*)-7-pentacosene (98%), (*Z*)-9-pentacosene (97%), (*Z*)-5-heptacosene (99%), (*Z*)-7-heptacosene (97%), (*Z*)-9-heptacosene (97%), (*Z*)-5-nonacosene (99%), (*Z*)-7-nonacosene (97%), and (*Z*)-9-nonacosene (97%), were synthesized following literature procedures (3). Briefly, terminal alkynes (1.0 eq) were deprotonated using 1M n-BuLi (1.05 eq) in hexane in dry THF/HMPA (2:1) at −70 °C for 30 min, then coupled to alkyl bromides (1.0 eq) at −70 °C to room temperature (rt) for 4 h. The alkynes were then reduced to *cis*-alkenes using H_2_ and Lindlar catalyst (5% Pd/BaSO_4_) in the presence of quinolene in hexane at rt. Heptacosane (99%) and nonacosane (99%) were prepared from further reduction of corresponding alkenes using hydrogen with 5% Pd/C catalyst in hexane at room temperature. All products were purified by vacuum removal of semi-volatile impurities and then silica gel chromatography, eluting with hexane.

##### GC-MS and GC-EAD

An HP7890-5975C (Agilent, Palo Alto, USA) GC-MS system was used with 1 mL/min helium as carrier gas. The splitless inlet with a small inner diameter liner for SPME samples (Supelco, Bellefonte PA, USA) was heated to 250 °C. An HP-5ms (30 m × 250 μm × 0.25 μm film, Agilent, Palo Alto, USA) column was used. The oven ramp was set as 50 °C for 1 min and then 10 °C/min to 280 °C for 10 min. The transfer line was 250 °C. In the quadrupole mass spectrometer, a 70 eV EI ion source was used, heated to 230 °C. The mass range scanned was *m/z* 28.5–500 at 4 scan/s. The abundance threshold for detection was set to 10. Data were collected and analysed using MSD Chemstation software (Agilent Technologies, Palo Alto CA, USA). Based on the literature (4,5), all compounds were identified by comparing their retention times and mass spectra to those of the synthetic standards.

For GC analyses, an HP7890B GC (Agilent) was used to separate the HS-SPME extracts with 1.0 mL/min helium as carrier gas. The splitless inlet was heated to 260 °C. An HP-5 (30 m × 320 μm × 0.32 μm, Agilent) column was used. The oven ramp was set as 50 °C for 1 min and then 10 °C/min to 280 °C for 10 min. FID signals were recorded and analysed using Chemstation software. Analyses of HS-SPME-GC were replicated 9 times (3 colonies each by 3 times).

I used a custom-made GC-EAD and EAG system (6,7) to test the antennal electrophysiological responses of undertaking bees. The cuticle with two antennal bases of each undertaking bee was dissected from the head capsule with a scalpel. The preparation was mounted to the reference electrode by surface tension of the Ringer’s solution^6^. The tips of the antennae were cut off using iris scissors under a microscope.

The tips of each of the two antennae were mounted to two LMP7721 (Texas Instruments, Dallas, TX, USA) based probes using glass electrodes filled with Ringer’s solution. Ultra-pure and clean platinum wires (0.4 mm) were used to connect the electrodes and input pin of the amplifier to prevent any electrolysis potential. The EAD signals of both antennae were fed to an HP 34465A digital multimeter (Keysight, Santa Rosa, USA) in DC measuring mode at 10 reads per second. Signal data were recorded using BenchVue software (Keysight). In GC-EAD analyses, an HP7890B GC (Agilent) was used to analyse the CHC solvent extracts with 1.0 mL/min helium as carrier gas. The splitless injector was heated to 280 °C. An HP-5 (30 m × 320 μm × 0.32 μm film, Agilent) column was used. The oven ramp was set at 200 °C for 1 min and then 10 °C/min to 280 °C for 10 min. The EAD transfer line was heated to 280 °C to prevent condensation of CHCs. The column effluents were diluted by a 42 cm/s clean, humidified, static-free (filter by grounded copper mesh) air flow in a PTFE odour pipette (8.0 mm in diameter) before being delivered to the antennal preparation. One bee equivalent of CHCs in 2 μL hexane was injected for each test. The GC-EAD tests were repeated 9 times. Only reproducible responses in all samples were considered EAD active responses.

##### CHC vapour pressure calculations

Vapour pressures (*VP*) of CHCs were calculated using the MPBPVP module in EPI Suite 4.11 (EPA, New York, USA). Three methods were used for calculation, the Antoine method (8), Modified Grain method^9^, and Mackay method (9). The ratio of *VP*s at high and low temperatures, including *VP*_45°C_/*VP*_15°C_ and *VP*_45°C_/*VP*_25°C_ were calculated and plotted for comparison. For consistency, no experimental data were used in calculation.

#### Statistical analysis

The effects of samples, colonies, and environment temperature on the counts of samples removed/tested were analysed using a Generalized Linear Mixed Model (GLMM) with Binary Probit (for dissections test and heated and/or washed dead bee test) or Poisson Loglinear (for wax concentration test) as link function. Estimated marginal means were compared using the Sequential Bonferroni method. Mean temperatures of active living bees were estimated from the sampled bees using a *t-test*. The differences of compound quantities in dead and living bees were compared using a paired-sample *t-test*. In the synthetic CHC assays, the effect of sample, colony, and environmental temperature on the times of dead bee removal were analysed using a Generalized Linear Model (GLM). The estimated marginal means (EM: mean±error) were compared using the Sequential Bonferroni method.

## Supplementary Figures and Table

**Figure S1.**
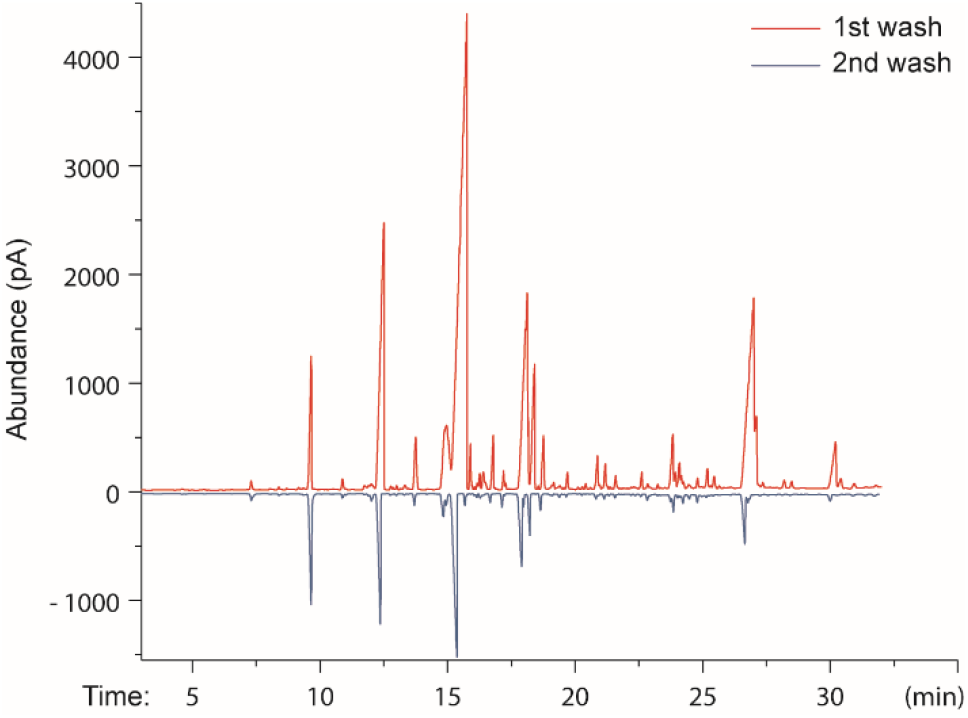
Solvent wash reduced not eliminated the CHC

**Figure S2.**
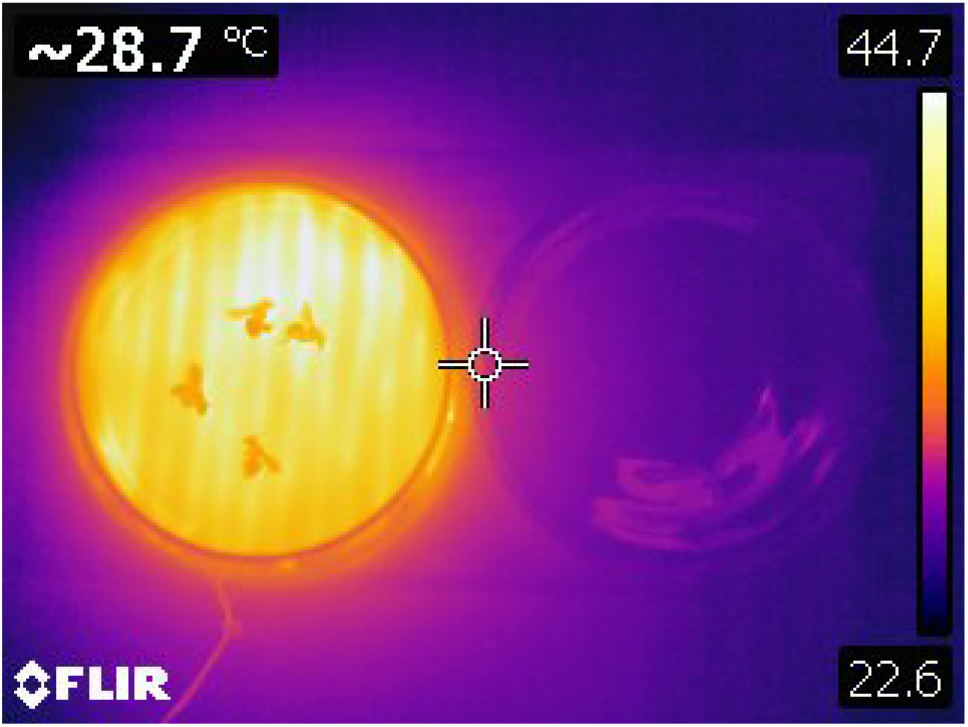
Thermo image of the heating apparatus

**Figure S3.**
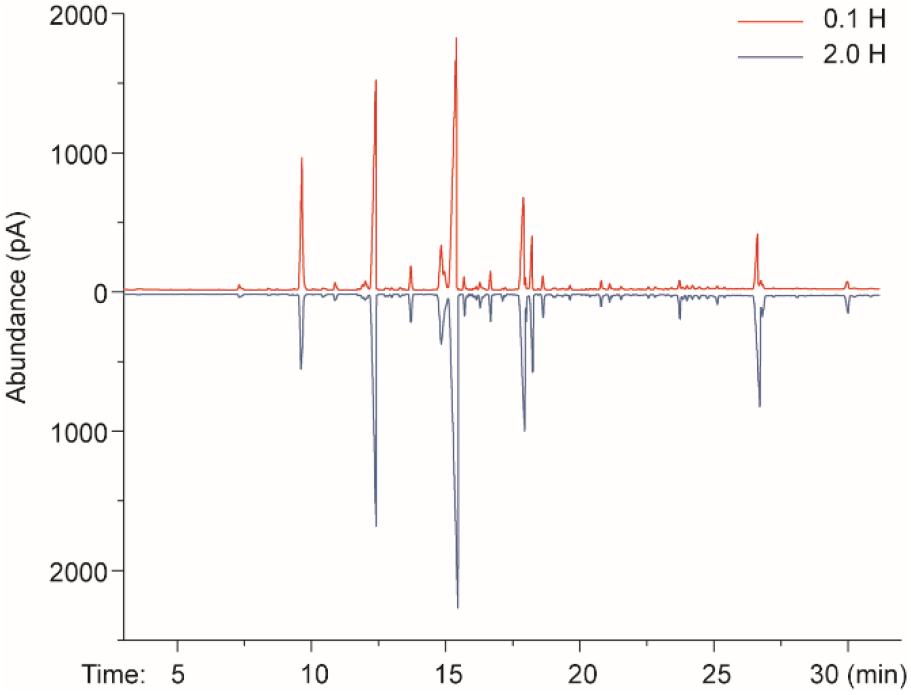
CHC residue did not change fast with time after death

**Figure S4.**
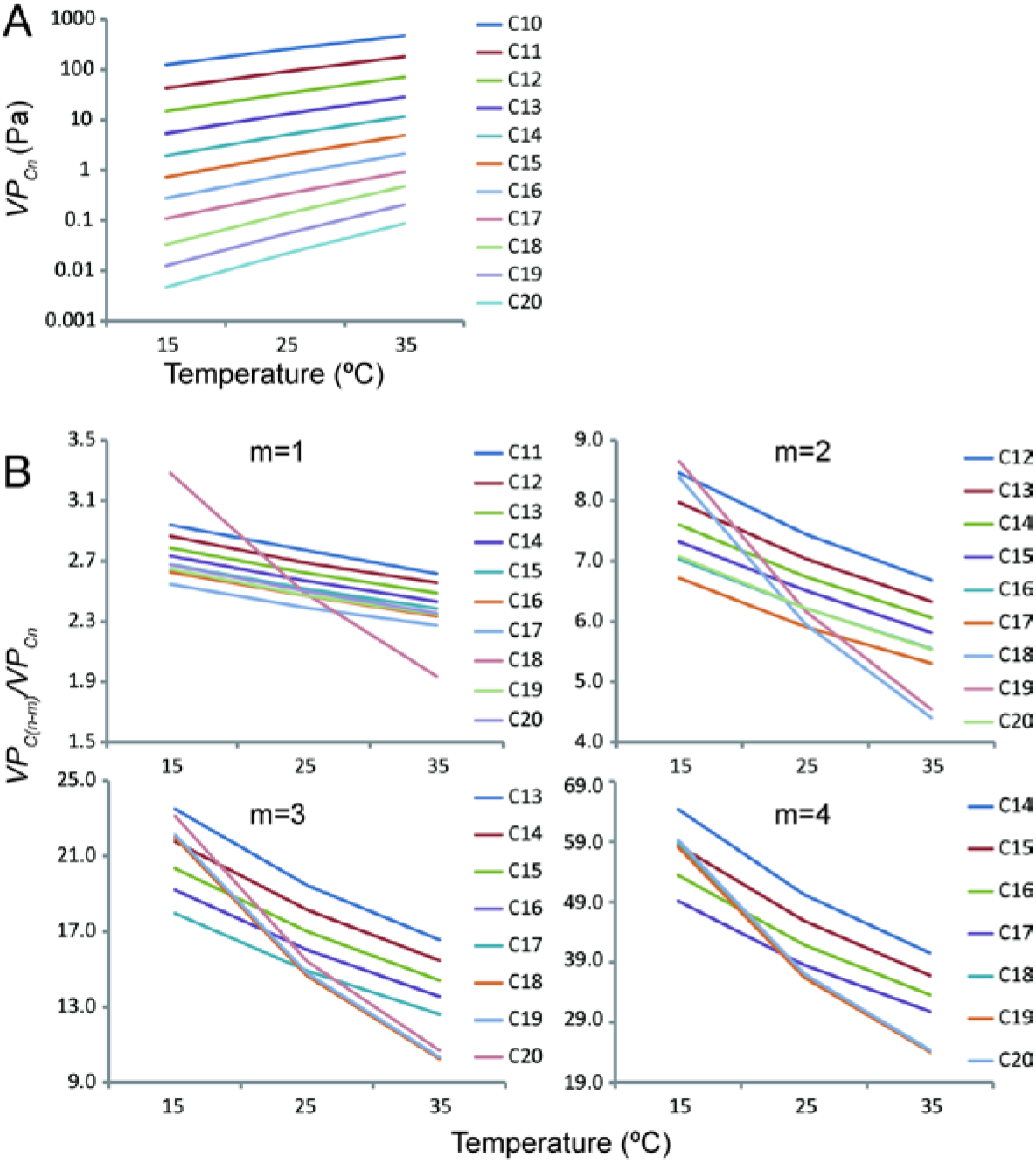
Temperature change the ratio of hydrocarbons vapour pressure. A Calculated vapour pressure of C10-C20 n-alkanes using Antoine method. B Changed *VP* ratio between the alkanes with 1, 2, 3, 4 carbon differences in molecule sizes.

**Table S1.** Sizes of tested colonies.

| Colony No. | Number of individuals |
| --- | --- |
| 1 | 500~1000 |
| 2 | 3000~3500 |
| 3 | 2000~3000 |
| 4 | 3500~4000 |
| 5 | 5000~6000 |

